# A method for the efficient iron-labeling of patient-derived xenograft cells and cellular imaging validation

**DOI:** 10.1101/2021.01.12.426411

**Authors:** Natasha N. Knier, Veronica P. Dubois, John A. Ronald, Paula J. Foster

**Affiliations:** Department of Medical Biophysics, Western University, London, Ontario Canada; Imaging Laboratories, Robarts Research Institute, London, Ontario, Canada; Lawson Health Research Institute, London, Ontario, Canada

## Abstract

There is momentum towards implementing patient-derived xenograft models (PDX) in cancer research to reflect the histopathology, tumour behavior, and metastatic properties observed in the original tumour. These models are more predictive of clinical outcomes and are superior to cell lines for preclinical drug evaluation and therapeutic strategies. To study PDX cells preclinically, we used both bioluminescence imaging (BLI) to evaluate cell viability and magnetic particle imaging (MPI), an emerging imaging technology to allow for detection and quantification of iron nanoparticles. The goal of this study was to develop the first successful iron labeling method of breast cancer cells derived from patient brain metastases and validate this method with imaging during tumour development.

Luciferase expressing human breast cancer PDX cells (F2-7) were successfully labeled after incubation with micron-sized iron oxide particles (MPIO; 25 μg Fe/mL). NOD/SCID/ILIIrg^-/-^ (n=5) mice received injections of 1×10^6^ iron-labeled F2-7 cells into the fourth mammary fat pad (MFP). BLI was performed longitudinally to day 49 and MPI was performed up to day 28. *In vivo* BLI revealed that signal increased over time with tumour development. MPI revealed decreasing signal in the tumours and increasing signal in the liver region over time.

Here, we demonstrate the first application of MPI to monitor the growth of a PDX MFP tumour. To accomplish this, we also demonstrate the first successful labeling of PDX cells with iron oxide particles. Imaging of PDX cells provides a powerful system to better develop personalized therapies targeting breast cancer brain metastasis.

## Introduction

Breast cancer is one of the most common cancers seen in women, currently affecting 1 in 8 women in North America (1). Mortality associated with this disease is caused most frequently by metastasis, which is the spread of cancer from the primary tumour to other distant locations in the body (2). In breast cancer, these locations often include the brain, bone, lungs, and lymph nodes (3). For brain metastases in particular, prognosis is poor, with mean 1-year survival rates of only 20% (4), and 2-year survival rates of <2% (5). Additionally, the incidence of brain metastasis (BM) is increasing, as neuroimaging techniques improve and treatments that allow for longer patient survival permits more time for cells to metastasize to the central nervous system (6,7). Insight into the mechanisms and pathophysiology of breast cancer brain metastasis have long relied on the use of immortalized cell lines that have been injected intracardially into the left ventricle of the mouse heart in order to selectively grow tumours within the brain. While these cell lines have been well characterized, they do not represent the tumour heterogeneity, metastatic behaviours, and histopathology seen clinically, and are unsuitable for evaluating therapies due to their fast progression *in vivo* (8,9).

Patient-derived xenografts, or PDXs, have begun to supplant traditional cell lines due to their retention of clinical biomarkers and heterogeneity from the original tumour (10). In recent years, PDXs have been developed to grow in the mammary fat pad of NOD/SCID/ILIIrg^-/-^ (NSG) mice, with a long-term objective to advance personalized medicine. This strategy has shown exciting progress for the development of novel PDXs from brain metastases in breast cancer patients. In 2017, Contreras-Zárate et al. developed BM-PDXs to study the biology of brain metastasis and to serve as tools for testing novel therapeutic approaches (11). These novel BM-PDXs retained intratumoural heterogeneity and metastatic potential, providing a clinically relevant model to study mechanisms of brain metastatic colonization and slower progression to allow for therapeutic testing. Currently, most PDX models are typically studied using methods such as histology, immunohistochemistry, and fluorescent microscopy, limiting the ability to study these models before an experimental endpoint has been reached. Tools that enable the longitudinal study of the fate of BM-PDXs would provide valuable information to characterize to the mechanisms and metastatic patterns *in vivo*.

Cellular imaging and cell tracking can be used to study cancer cell populations and metastatic processes *in vivo*. Bioluminescence imaging (BLI) with the reporter firefly luciferase (Fluc) has been widely utilized for tracking preclinical models of cancer. BLI is a valuable imaging modality as it allows for the longitudinal study of cell viability. Fluc BLI requires adenosine triphosphate (ATP) as a co-factor, and thus, the signal measured with BLI is directly proportional to the number of viable cells in a region of interest (12). This modality is exceptionally useful when measuring treatment and therapeutic response, as cell viability may change but tumour volumes can remain unaltered.

Our group has previously shown that BLI can be complemented with iron-based cellular MRI technology to provide longitudinal measures of cancer cell viability in preclinical models (13,14). Iron-based cellular MRI requires cells to be loaded with superparamagnetic iron oxide nanoparticles (SPIONs) and has shown to provide single cell sensitivity (15). However, a limitation of this modality is that SPIONs create regions signal loss where iron-labeled cells are present in images, and so, determining the quantitation of signal loss is challenging and it is not possible to determine cell number (16). Magnetic particle imaging (MPI) is an emerging imaging modality that directly detects SPIONs (17,18), resulting in a positive signal that appears as a “hot spot” in images. The signal strength detected is linearly proportional to the number of SPIONs, allowing for quantitation (19). Presently, MPI has been used as a cell tracking modality for immortalized cancer cell lines (20,21), stem cells (22–25), pancreatic islets (26), T-cells (27) and macrophages (28,29), however, no studies exist studying the growth of patient-derived xenografts labeled efficiently with an SPION *in vivo*.

Efficient iron labeling of a patient-derived xenograft cell line presents a challenge due to the mixed and heterogeneous population of cells. In this work, we report the first iron-labeling method for a patient-derived xenograft cell line and validate its utility for cell tracking with MPI and BLI.

## Matesrials and Methods

### F2-7 Cell Line Origin and Cell Culture

A firefly luciferase and enhanced green fluorescent protein (eGFP) expressing patient-derived xenograft cell line (F2-7) was obtained from the Cittely Lab (UC Denver), and was originally developed from a triple-negative brain metastatic patient-derived xenograft (11). F2-7 cells were maintained in T75cm^2^ flasks coated with collagen-I (1mg/mL) for 2 hours to encourage attachment. 12mL of Dulbecco’s modified Eagle’s medium (DMEM)-F12 supplemented with 10% of fetal bovine serum (FBS), 1 μg/ml hydrocortisone, 100 ng/ml of cholera toxin, and 1 nM of insulin was added to flasks after collagen coating. Cells grew under 37°C and 5% CO_2_. Since cells grow as both non-adherent mammospheres and adherent single cells, both populations require maintenance with each passage.

### MPIO Labeling Procedure

#### Day 1

DMEM-F12, PBS, and trypsin were heated at 37°C for 30 minutes in a water bath. Conditioned media and non-adherent cells were collected into 15mL conical tubes from confluent flasks of F2-7 cells. Conical tubes were centrifuged with a ThermoFisher Cytospin 4 for 5 minutes at 900 rpm, and the supernatant was removed and stored for future use as conditioned media. 1mL of 0.25% Trypsin-EDTA was added to the cell pellet within the conical tube and was placed in the water bath for 2 minutes to trypsinize cells. In parallel, 1mL of 0.25% Trypsin-EDTA was added to the flask with remaining adherent cells and incubated for approximately 5 minutes. Once adherent cells and mammospheres were dissociated, cell populations were mixed and resuspended in 5mL of fresh media. After, the mixed population of cells were centrifuged again at 900rpm for 5 minutes. The supernatant with trypsin was removed, and the cell pellet was resuspended in 2-5mL. Cells were assessed for viability with Trypan blue exclusion assay and counted to achieve the correct volume of media to plate 2×10^6^ cells. Cells were plated in uncoated T75cm^2^ flasks to discourage attachment with 50% fresh media and 50% conditioned media. To iron label these cells, 3 methods of iron labeling were performed to determine the most efficient labeling technique. In the first trial, cells were supplemented with 25 μg Fe/mL of micron-sized iron oxide particles (MPIO) (0.9 μm diameter, ~63% magnetite, labeled with Flash Red; Bangs Laboratory, Fishers, IN, USA), and swirled in the flask to mix. The second trial used the 50 μg Fe/mL of the same MPIO, and the third used 25 μg Fe/mL with a magnetic plate under the flask to allow for magnetofection (30). Following this, all flasks were incubated for 24 hours.

#### Day 2

After 24 hours, cells from the flasks were harvested and dissociated in accordance to the protocol in Day 1. After, cells were harvested and washed with 10mL PBS in the flask to remove unincorporated MPIO. Cells were centrifuged at 900rpm for 5 minutes. This process was repeated thrice to thoroughly remove unincorporated MPIO prior to cell injections. Cell viability was assessed and calculated using the Trypan blue exclusion assay. A simplified visual of this protocol is shown in Figure 1.

**Figure 1.**
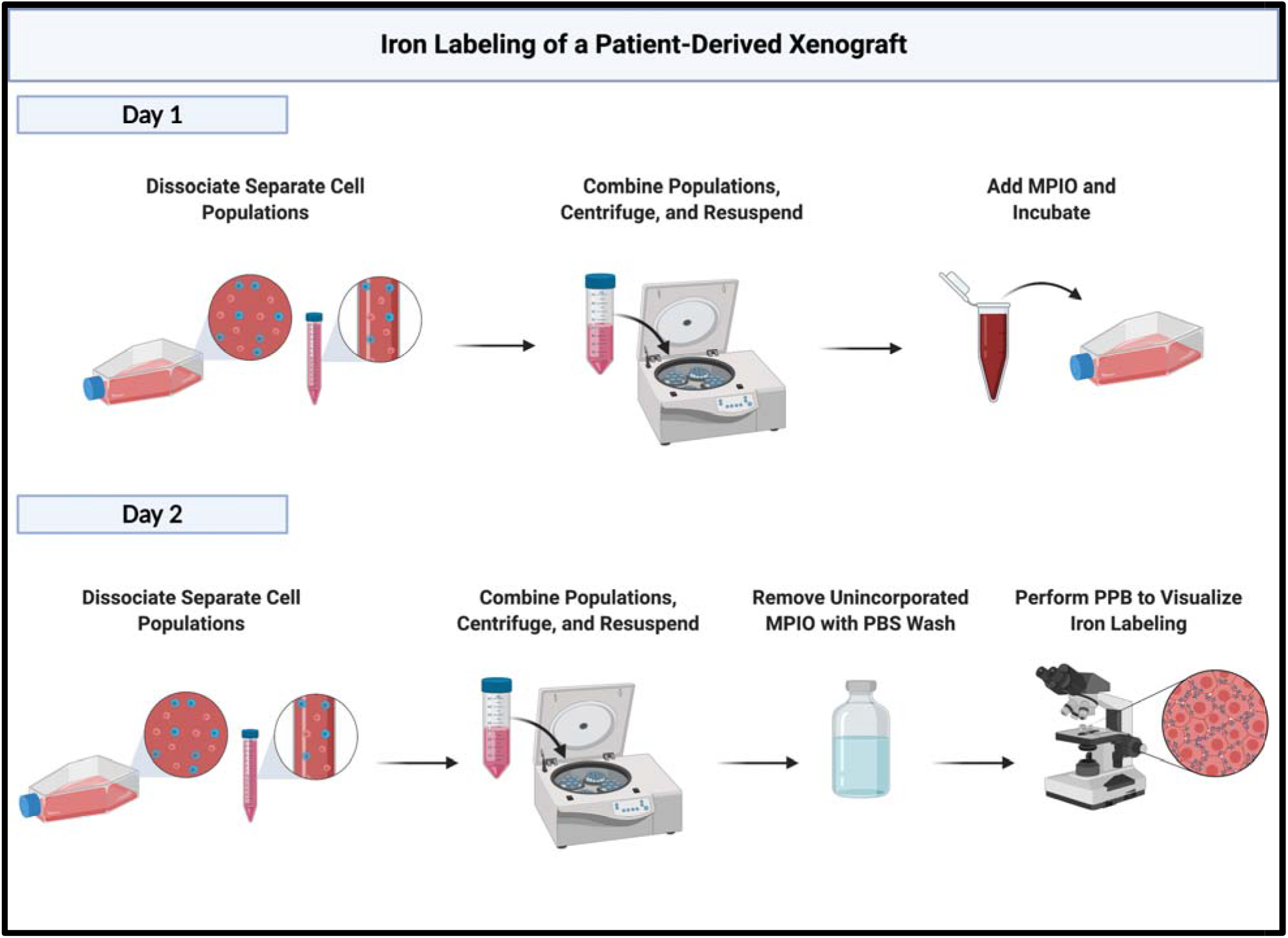
A simplified illustration of the workflow to efficiently label patient-derived xenograft cells with iron-oxide particles. Created with BioRender.com.

### Assessing Iron Labeling

To visualize MPIO labeling, labeled cells were affixed to a glass slide with a ThermoFisher Cytospin 4 cytocentrifuge and fixed with a Methanol/Acetic acid solution. Slides were then stained with a Perl’s Prussian Blue (PPB) solution and counterstained with Nuclear Fast Red. Slides were dehydrated with increasing concentrations of ethanol, cleared with xylene, and coverslipped with a xylene-based mounting medium. These PPB-stained slides were examined to assess the localization of MPIO within the cell to determine the labeling efficiency using a Zeiss AXIO Imager A1 Microscope (Zeiss Canada, Toronto, ON, Canada). Iron oxide particles appear dark blue, and cells appear light pink in colour.

### Animal Model and Workflow

All animals were cared for in accordance with the standards of the Canadian Council on Animal Care, under an approved protocol of the University of Western Ontario’s Council on Animal Care and housed in a pathogen-free barrier facility. Female NSG mice (n=5) (NOD.Cg-PrkdcscidIl2rgtm1Wjl/SzJ, 6-8 weeks, Humanized Mouse and Xenotransplantation Facility, Robarts Research Institute, University of Western Ontario, London, ON) were first anesthetized with isoflurane (2% in 100% oxygen). NSG mice were then injected into the fourth mammary fat pad (MFP) with a suspension of 1×10^6^ MPIO-labeled F2-7luc/eGFP+ cells in 50μL of sterile saline and 50uL of Matrigel. Mice were imaged with BLI (n=2) out to day 49 and MPI (n=3) out to day 28. At endpoint, mice were sacrificed by isoflurane overdose. Tumours were excised and placed in paraformaldehyde for an additional 24 hours. *Ex vivo* MFP tumour volumes were estimated using the following formula = 0.52 (width)^2^ (length) to approximate the volume of an ellipsoid (mm^3^).

### *In Vitro* BLI Procedure

To evaluate the relationship between cell number and BLI signal, cells were seeded in 24-well plates in 0.5mL of growth medium at concentrations of 2×10^5^, 6×10^5^, 1×10^6^ cells per well. Cells were allowed to adhere for 24 hours and then 5μL of D-luciferin (30mg/mL) was added to the cell media 2 minutes prior to imaging using a hybrid optical/X-ray scanner (IVIS Lumina XRMS In Vivo Imaging System, PerkinElmer). Region-of-interest (ROI) analysis was performed for each well using Living Image Software (IVIS Imaging Systems, PerkinElmer) and data is expressed as the average radiance (photons/sec/cm^2^/steradian).

### *In Vivo* BLI Procedure

BLI was performed on NSG mice (n=2) on days 4, 28, 35, 42, and 49 using a hybrid optical/X-ray scanner (IVIS Lumina XRMS In Vivo Imaging System, PerkinElmer). Mice were anesthetized with isoflurane (2% in 100% oxygen) using a nose cone attached to an activated carbon charcoal filter. Mice received a 100uL intraperitoneal injection of D-luciferin (30mg/mL), and luminescent images were captured for approximately 35 minutes to obtain the peak signal at each imaging session. Region-of-interest (ROI) analysis was performed for each mouse using Living Image Software (IVIS Imaging Systems, PerkinElmer) and data is expressed as the average radiance (photons/sec/cm^2^/steradian).

### MPI Acquisition

Full body MPI images of mammary fat pad tumour-bearing NSG mice (n=3) were acquired at days 0 (30 minutes post-injection), 7, 14, and 28. Images were collected on a Momentum™ scanner (Magnetic Insight Inc., Alameda, CA, USA) using the 3D high sensitivity isotropic (multichannel) scan mode. In this mode, images were acquired using a 5.7 T/m gradient, 21 projections and a FOV (field of view) of 12 x 6 x 6 cm, for a total scan time ~18 mins per mouse. Mice were anesthetized with 2% isoflurane in 100% oxygen for the entirety of the scan. 3D high sensitivity isotropic images of *ex vivo* tumours (n=2) removed on Day 40 were acquired using the same parameters and a FOV of 4 x 6 x 6 cm, for a total scan time of ~12 minutes.

### MPI Calibration and Signal Quantification

To generate a calibration curve for converting MPI signal to iron content, samples were made with 2 μL aliquots of MPIO and 2 μL PBS and were imaged using the 3D high sensitivity isotropic parameters. The FOV was 12 x 6 x 6 cm. The following samples were scanned separately: 0.175 μg, 0.35 μg, 0.70 μg, 1.40 μg, 2.80 μg, 5.6 μg. Images were analyzed using the open-source Horos imaging software, version 3.3.5 (Annapolis, MD USA). To quantify the MPI signal in each image set, signal intensities were set to full dynamic range in order to represent the full range of signal in each specific ROI, such as the calibration samples, mammary fat pad tumours, and liver region. Areas of interest were then segmented manually, and 3D volumes were reconstructed and calculated using the Horos volume algorithm. The total MPI signal was calculated using the equation *mean signal x volume*(*mm*^3^).

### Statistical Analysis

Statistical analysis was performed using GraphPad Prism 8 Software (GraphPad, San Diego, CA, SA). Pearson’s rank correlation was used to determine the relationship between total MPI signal and iron content. *In vivo* data was expressed as mean ± SD and analyzed with a one-way ANOVA. Differences were considered statistically significant at p < 0.05.

## Results

### Cell Labeling

For each trial of labeling, F2-7 cells had varying labeling efficiencies demonstrated by a PPB stain shown in Figure 2, with cancer cells appearing pink, and intracellular iron in blue. For trial 1 with 25 μg Fe/mL of MPIO, F2-7 cells were efficiently labeled with labeling efficiency of 81.80±10.14% (Fig. 1A) and viability of over 90%. This labeling efficiency was deemed successful and the 25 μg Fe/mL of MPIO labeling procedure was used for the remainder of the study. The second trial, using the 50 μg Fe/mL of the same MPIO, resulted in a labeling efficiency of 27.51±1.19% (Fig. 1B). Finally, the third trial which used 25 μg Fe/mL with a magnetic plate resulted in a labeling efficiency of 5.55±1.65% (Fig. 1C).

**Figure 2.**
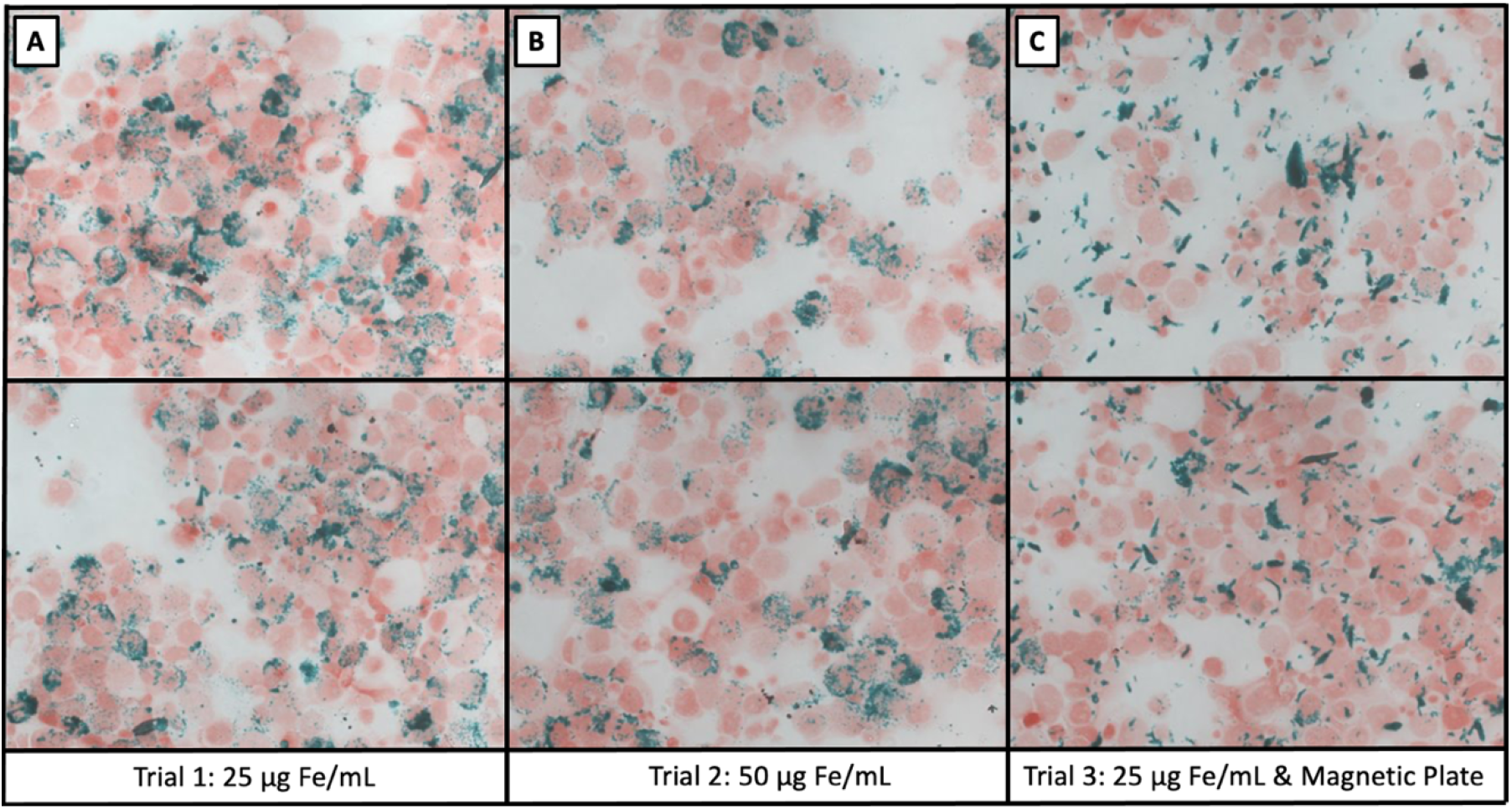
**A)** Perl’s Prussian Blue (PPB) stain showing F2-7 cells labeled with 25 μg Fe/mL of MPIO, with labeling efficiency of 81.80±10.14%. **B)** PPB stain showing F2-7 cells labeled with 50 μg Fe/mL of MPIO, with a labeling efficiency of 27.51±1.19%. **C)** PPB stain showing F2-7 cells labeled with 25 μg Fe/mL and a magnetic plate, resulting in a labeling efficiency of 5.55±1.65%.

### Bioluminescence Imaging (BLI)

F2-7/eGFP-luc cells were seeded at concentrations of 2×10^5^, 6×10^5^, 1×10^6^ cells per well and *in vitro* BLI was performed (Fig. 3A). A significant positive correlation was found between t**he** number of F2-7/eGFP-luc cells and BLI signal (R^2^ = 0.9664). Specifically, as seeded cell number increased, BLI signal also increased (Fig. 3B). In Figure 3C the BLI signal from a representative tumour is shown to increase over time with tumour development. No BLI signal was detected in any other region of the mouse body.

**Figure 3.**
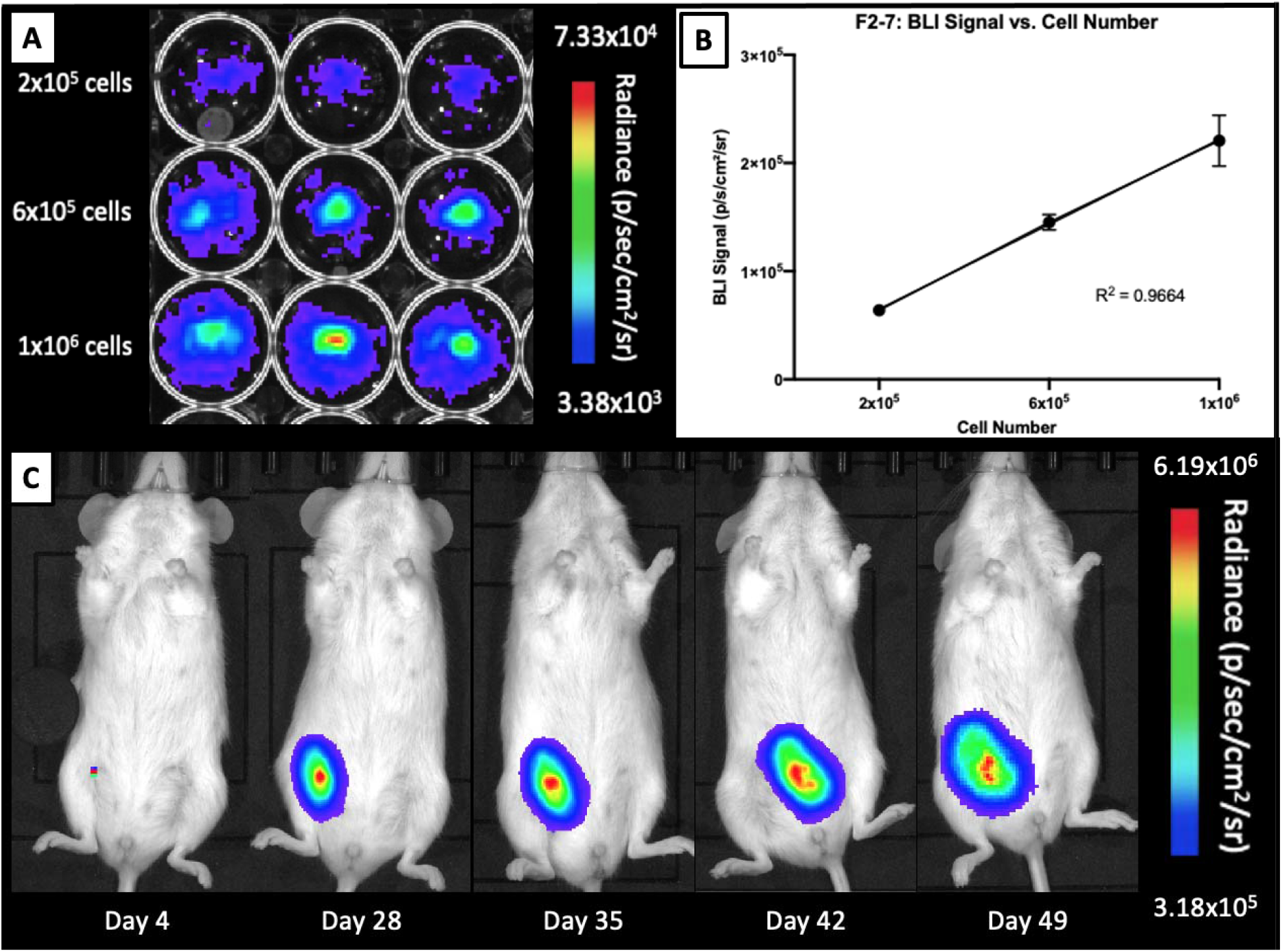
**A)** F2-7luc/eGFP+ cells seeded at various concentrations. **B)** A strong linear correlation is seen between cell number and BLI signal; R^2^□=□0.9664. **C)** BLI signal from a representative tumour is shown to increase over time with tumour development from day 4 to day 49.

### Magnetic Particle Imaging (MPI)

A calibration line was generated to determine iron content for a given MPI signal based on the 3D, high sensitivity, isotropic parameters used to image MPIO. An example of the images of samples measured to generate calibration curves are shown in Figure 4A for MPIO and the calibration line generated from this data is shown in 4b. Based on this data, we determined here was a strong linear relationship between iron content and MPI signal (arbitrary units, A.U.) for MPIO (R^2^ = 0.9836, p <.001). The equation of the line was: MPI Signal = 69.559 (Iron Content) for MPIO. Using this relationship, iron content could be determined for a given MPI signal.

**Figure 4.**
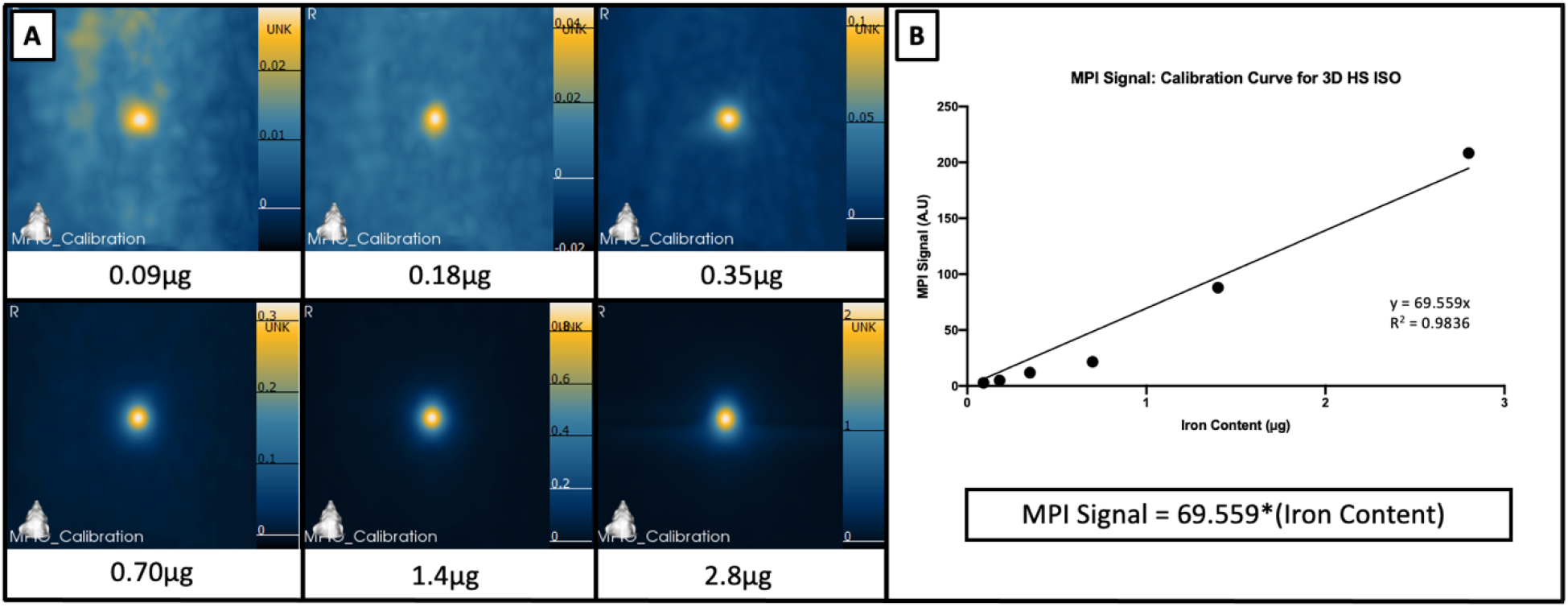
**A)** Images of MPIO samples measured to generate calibration curves. **B)** A strong linear relationship is seen between iron content and MPI signal; R2 = 0.9836

Figure 5A shows representative MPI of a tumour bearing mouse on day 0, 7, 14, and 28 postinjection of 1×10^6^ MPIO-labeled F2-7 cells. MPI signal is clearly visualized in the lower, right MFP tumour, with signal decreasing over time. Signal can also be seen in the liver region and is shown to increase in signal over time (Fig. 5B). The mean iron content in the MFP tumours decreased significantly between day 0 (M=4.06 ± 2.09 μg) and day 28 (M= 0.41 ± 0.25 μg) (p=0.0095). Conversely, the mean iron content measured from the MPI signal in the liver region was 2.66 ± 0.94 μg on day 0 and 3.38 ± 1.16 μg on day 28 (Fig. 5C). Additionally, *ex vivo* MPI was performed on tumours removed on day 40 (Fig. 5D). A representative *ex vivo* tumour is shown in Figure 5E. The tumours measured 152.88 mm^3^ and 178.36 mm^3^, and the iron content was 0.05 μg and 0.75 μg, respectively.

**Figure 5.**
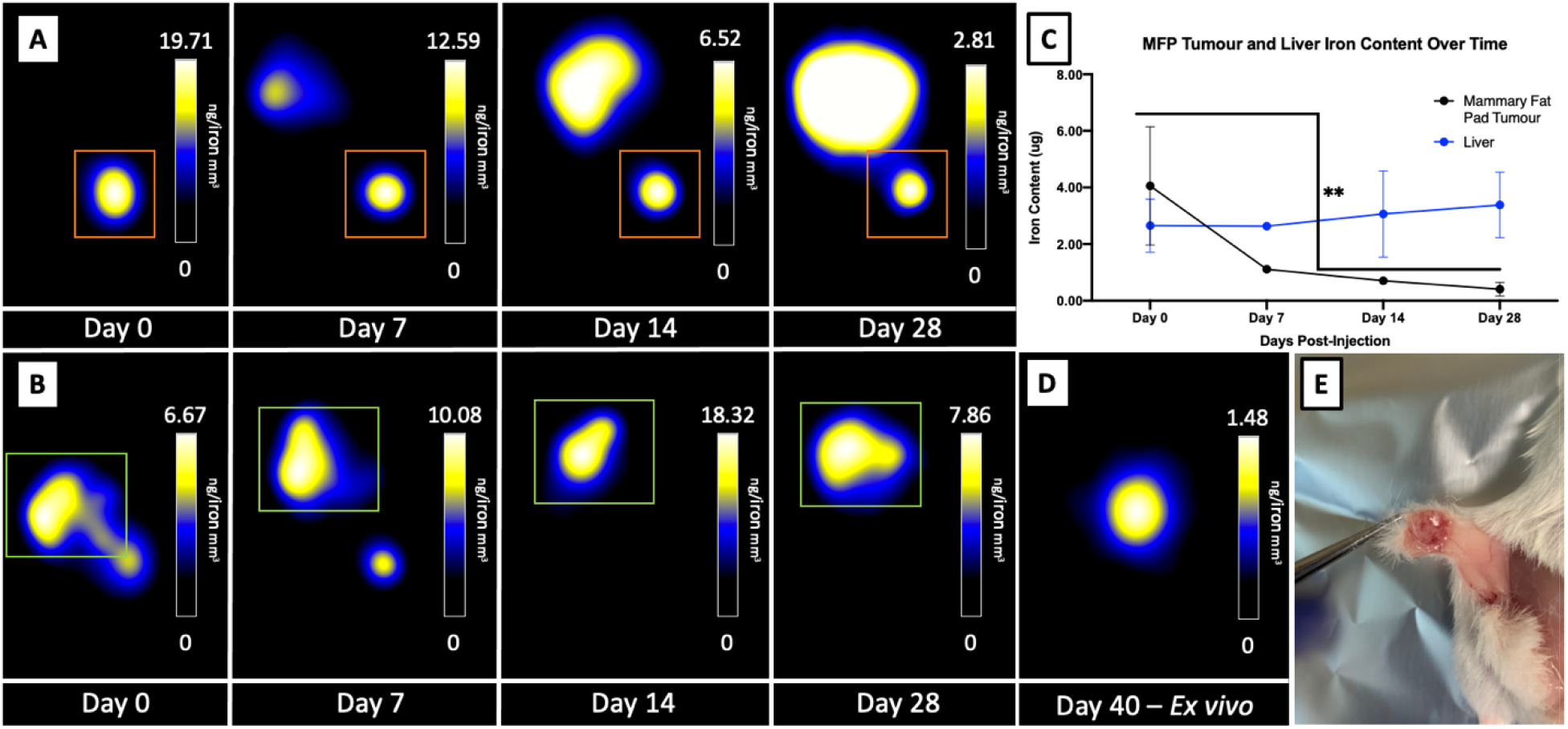
**A)** Representative MPI of a tumour bearing mouse on day 0, 7, 14, and 28 post-injection of 1×10^6^ MPIO-labeled F2-7 cells, with images window-levelled to the tumour (orange box). **B)** Representative MPI of a tumour bearing mouse on day 0, 7, 14, and 28 post-injection of 1×10^6^ MPIO-labeled F2-7 cells, with images window-levelled to the liver (green box). **C)** Quantification of mean iron content over time in MFP tumours decreasing significantly between day 0 (M=4.06 ± 2.09 μg) and day 28 (M= 0.41 ± 0.25 μg) (p=0.0095), and the liver region on day 0 (2.66 ± 0.94 μg) and on day 28 (3.38 ± 1.16 μg). **D)** Representative MPI of an ex vivo tumour removed on day 40. **E)** A representative ex vivo tumour of F2-7 cells.

## Discussion

The increasing incidence of breast cancer brain metastases and its poor prognosis has highlighted the critical need for clinically relevant models to develop new therapeutic strategies and to understand the mechanisms underlying its progression. PDX models have been used increasingly to understand the role of tumour heterogeneity in the development of novel drugs and metastatic progression, and while other groups have made exciting progress in this area, most methods employed have been performed *ex vivo*, through histology, or with immunohistochemistry. Alone, these do not allow for the longitudinal study of cancer progression *in vivo*, and therefore, tools such as cellular imaging would be extremely valuable in further investigation and characterization of these models. Additionally, only a handful of studies exist that validate the engraftment and growth patterns of PDX models with imaging. At this time, only three studies have used MRI to image PDX models of breast cancer brain metastasis. In 2016, Ni et al. monitored therapeutic response to combination therapies by both MRI and BLI of an orthoptic PDX brain metastatic model (31). In 2017, two novel brain metastatic PDX cell lines, including F2-7, as well as BM-E22-1, were imaged to determine successful engraftment and detect metastases within the mouse brain (32). Sharma et al. also confirmed the tumour establishment of two PDX cell lines (PDX2147 and PDX1435) with MRI after 30 days (33). Including the mentioned study by Ni et al., only three studies have used BLI to monitor cell viability in PDX models of breast cancer brain metastasis. Turner et al. used BLI to monitor the response of two basal-like, triple-negative PDX cell lines (WHIM2 and WHIM30) to treatment (34). Liu et al. described establishment of orthotopic mouse models of BCBM-PDXs and monitored their engraftment with BLI (35). To address this, we developed a novel method to label luciferase expressing PDX cells with iron-oxide particles to allow for the *in vivo*, longitudinal cell tracking with two imaging modalities for the first time.

Our group has previously demonstrated that BLI is a valuable tool for cell tracking preclinical models of breast cancer brain metastasis, as is it is able to provide longitudinal measures of tumour growth and indicates cell viability (13). MPI is an emerging, highly sensitive imaging modality that can be used for cell tracking and offers the benefits of directly detecting iron-oxide nanoparticles to allow for the quantification of iron in a region of interest. While MPI has been used to detect cell lines of breast cancer (20,21,36), colon cancer (37), and tumour-associated macrophages (28), to date, no studies exist demonstrating the labeling of PDX cells with iron-oxide nanoparticles to allow for MPI cell tracking.

In this work, we visualized the iron labeling of the luciferase expressing F2-7-eGFP with PPB stains and demonstrated that 25 μg Fe/mL was the most successful labeling trial. The addition of the magnetic plate for labeling was likely ineffective due to the fact that the majority of these cells are in suspension during labeling, which differs from most studies which have labeled adherent cells. Previous work by our group has shown that this concentration of MPIO effectively labels immortalized cancer cells for the purposes of cell tracking (38), however, the methods of labeling described in this work varies from the traditional labeling protocol of adherent cells due to the differences in homogeneity and complex cell culture required to grow PDX cells. Future work will investigate labeling trials with different iron particles and whether this changes the visualization of these cells *in vivo* with MPI or MRI.

In this model, F2-7 mammary fat pad tumours were monitored out to day 42 with BLI. BLI signal increased over time, demonstrating that the PDX cells had successfully engrafted and were proliferating over time. In contrast, the MPI signal from the MFP decreased over time during the 28 day period. This may be related to the clearance of iron released from labeled cancer cells. This is supported, in part, by the observation of MPI signal in the liver region, which was observed unexpectedly, and may occur as cancer cells in the developing tumour die and the iron particles are cleared. Wang et al. have shown that when iron-labeled islets were transplanted under the kidney capsule MPI signal is only detected in the kidney on day 1 post-transplant but on day 14 MPI signal also appears in the liver, where the released iron particles accumulate (26). These observations require more study. Future work will investigate this potential transfer of iron from the MFP to the liver region through *ex vivo* imaging and iron quantification with MPI.

Our group and others have used either BLI or MPI to detect iron-labelled cancer cells *in vivo*, however, only one study has used both modalities (36), although they performed imaging at a single timepoint in a single mouse. We have previously shown that labeling cancer cells with iron-oxide nanoparticles does not significantly affect cell viability, proliferation, apoptosis, or metastatic efficiency, demonstrating that this labeling agent is an effective technique to track cells *in vivo* (15,39). With the increased use of PDX models as a platform to study cancer metastasis and develop novel drugs and therapeutics, this method provides a reliable and efficient method to determine the fate of these cells *in vivo* and their response to therapeutic treatments.

## Acknowledgements

The authors would like to thank the Canadian Institute for Health Research and the Breast Cancer Society of Canada for their funding.

## References

1. DeSantis C, Ma J, Bryan L, Jemal A. Breast cancer statistics, 2013. CA Cancer J Clin. 2014;64(1):52–62.

2. Chambers AF, Groom AC, MacDonald IC. Dissemination and growth of cancer cells in metastatic sites. Nat Rev Cancer. 2002;2(8):563–572.

3. Kennecke H, Yerushalmi R, Woods R, Cheang MCU, Voduc D, Speers CH, et al. Metastatic behavior of breast cancer subtypes. J Clin Oncol Off J Am Soc Clin Oncol. 2010 Jul 10;28(20):3271–7.

4. Shaffrey ME, Mut M, Asher AL, Burri SH, Chahlavi A, Chang SM, et al. Brain metastases. Curr Probl Surg. 2004 Aug 1;41(8):665–741.

5. Engel J, Eckel R, Aydemir Ü, Aydemir S, Kerr J, Schlesinger-Raab A, et al. Determinants and prognoses of locoregional and distant progression in breast cancer. Int J Radiat Oncol. 2003 Apr 1;55(5):1186–95.

6. Barnholtz-Sloan JS, Sloan AE, Davis FG, Vigneau FD, Lai P, Sawaya RE. Incidence Proportions of Brain Metastases in Patients Diagnosed (1973 to 2001) in the Metropolitan Detroit Cancer Surveillance System. J Clin Oncol. 2004 Jul 15;22(14):2865–72.

7. Fabi A, Felici A, Metro G, Mirri A, Bria E, Telera S, et al. Brain metastases from solid tumors: disease outcome according to type of treatment and therapeutic resources of the treating center. J Exp Clin Cancer Res CR. 2011 Jan 18;30(1):10.

8. Hidalgo M, Amant F, Biankin AV, Budinská E, Byrne AT, Caldas C, et al. Patient Derived Xenograft Models: An Emerging Platform for Translational Cancer Research. Cancer Discov. 2014 Sep;4(9):998–1013.

9. Gillet J-P, Calcagno AM, Varma S, Marino M, Green LJ, Vora MI, et al. Redefining the relevance of established cancer cell lines to the study of mechanisms of clinical anti-cancer drug resistance. Proc Natl Acad Sci. 2011 Nov 15;108(46):18708–13.

10. Dobrolecki LE, Airhart SD, Alferez DG, Aparicio S, Behbod F, Bentires-Alj M, et al. Patient-derived xenograft (PDX) models in basic and translational breast cancer research. Cancer Metastasis Rev. 2016;35(4):547–573.

11. Contreras-Zárate MJ, Ormond DR, Gillen AE, Hanna C, Day NL, Serkova NJ, et al. Development of novel patient-derived xenografts from breast cancer brain metastases. Front Oncol. 2017;7:252.

12. Prescher JA, Contag CH. Guided by the light: visualizing biomolecular processes in living animals with bioluminescence. Curr Opin Chem Biol. 2010 Feb;14(1):80–9.

13. Parkins KM, Hamilton AM, Makela AV, Chen Y, Foster PJ, Ronald JA. A multimodality imaging model to track viable breast cancer cells from single arrest to metastasis in the mouse brain. Sci Rep. 2016;6(1):1–9.

14. Hamilton AM, Parkins KM, Murrell DH, Ronald JA, Foster PJ. Investigating the impact of a primary tumor on metastasis and dormancy using MRI: new insights into the mechanism of concomitant tumor resistance. Tomography. 2016;2(2):79.

15. Heyn C, Ronald JA, Ramadan SS, Snir JA, Barry AM, MacKenzie LT, et al. In vivo MRI of cancer cell fate at the single-cell level in a mouse model of breast cancer metastasis to the brain. Magn Reson Med Off J Int Soc Magn Reson Med. 2006;56(5):1001–1010.

16. Makela AV, Murrell DH, Parkins KM, Kara J, Gaudet JM, Foster PJ. Cellular imaging with MRI. Top Magn Reson Imaging. 2016;25(5):177–186.

17. Bulte JWM. Superparamagnetic iron oxides as MPI tracers: A primer and review of early applications. Adv Drug Deliv Rev. 2019 01;138:293–301.

18. Zheng B, Yu E, Orendorff R, Lu K, Konkle JJ, Tay ZW, et al. Seeing SPIOs Directly In Vivo with Magnetic Particle Imaging. Mol Imaging Biol. 2017;19(3):385–90.

19. Wu LC, Zhang Y, Steinberg G, Qu H, Huang S, Cheng M, et al. A Review of Magnetic Particle Imaging and Perspectives on Neuroimaging. Am J Neuroradiol. 2019 Feb 1;40(2):206–12.

20. Melo KP, Makela AV, Hamilton AM, Foster PJ. Development of Magnetic Particle Imaging (MPI) for Cell Tracking and Detection. bioRxiv. 2020 Jul 13;2020.07.12.197780.

21. Parkins KM, Melo KP, Ronald JA, Foster PJ. Visualizing tumour self-homing with magnetic particle imaging. bioRxiv. 2020 Feb 19;2020.02.17.953232.

22. Nejadnik H, Pandit P, Lenkov O, Lahiji AP, Yerneni K, Daldrup-Link HE. Ferumoxytol Can Be Used for Quantitative Magnetic Particle Imaging of Transplanted Stem Cells. Mol Imaging Biol. 2019;21(3):465–72.

23. Zheng B, von See MP, Yu E, Gunel B, Lu K, Vazin T, et al. Quantitative Magnetic Particle Imaging Monitors the Transplantation, Biodistribution, and Clearance of Stem Cells In Vivo. Theranostics. 2016;6(3):291–301.

24. Bulte JWM, Walczak P, Janowski M, Krishnan KM, Arami H, Halkola A, et al. Quantitative “Hot Spot” Imaging of Transplanted Stem Cells using Superparamagnetic Tracers and Magnetic Particle Imaging (MPI). Tomogr Ann Arbor Mich. 2015 Dec;1(2):91–7.

25. Sehl OC, Makela AV, Hamilton AM, Foster PJ. Trimodal Cell Tracking In Vivo: Combining Iron- and Fluorine-Based Magnetic Resonance Imaging with Magnetic Particle Imaging to Monitor the Delivery of Mesenchymal Stem Cells and the Ensuing Inflammation. Tomography. 2019 Dec;5(4):367–76.

26. Wang P, Goodwill PW, Pandit P, Gaudet J, Ross A, Wang J, et al. Magnetic particle imaging of islet transplantation in the liver and under the kidney capsule in mouse models. Quant Imaging Med Surg. 2018 Mar;8(2):114–22.

27. Rivera-Rodriguez A, Hoang-Minh LB, Chiu-Lam A, Sarna N, Marrero-Morales L, Mitchell DA, et al. Tracking Adoptive T Cell Therapy Using Magnetic Particle Imaging. bioRxiv. 2020 Jun 3;2020.06.02.128587.

28. Makela AV, Gaudet JM, Schott MA, Sehl OC, Contag CH, Foster PJ. Magnetic Particle Imaging of Macrophages Associated with Cancer: Filling the Voids Left by Iron-Based Magnetic Resonance Imaging. Mol Imaging Biol. 2020 Aug 1;22(4):958–68.

29. Gaudet J, Mansfield J, Goodwill P. Imaging Cancer Immunology: Tracking Immune Cells in vivo with Magnetic Particle Imaging. J Immunol. 2019 May 1;202(1 Supplement):130.7–130.7.

30. Arbab AS, Yocum GT, Kalish H, Jordan EK, Anderson SA, Khakoo AY, et al. Efficient magnetic cell labeling with protamine sulfate complexed to ferumoxides for cellular MRI. Blood. 2004 Aug 15;104(4):1217–23.

31. Ni J, Ramkissoon SH, Xie S, Goel S, Stover DG, Guo H, et al. Combination inhibition of PI3K and mTORC1 yields durable remissions in mice bearing orthotopic patient-derived xenografts of HER2-positive breast cancer brain metastases. Nat Med. 2016 Jul;22(7):723–6.

32. Contreras-Zárate MJ, Ormond DR, Gillen AE, Hanna C, Day NL, Serkova NJ, et al. Development of Novel Patient-Derived Xenografts from Breast Cancer Brain Metastases. Front Oncol [Internet]. 2017 Nov 2 [cited 2020 Jul 27];7. Available from: https://www.ncbi.nlm.nih.gov/pmc/articles/PMC5673842/

33. Sharma S, Wu S-Y, Jimenez H, Xing F, Zhu D, Liu Y, et al. Ca2+ and CACNA1H mediate targeted suppression of breast cancer brain metastasis by AM RF EMF. EBioMedicine. 2019 Jun 1;44:194–208.

34. Turner TH, Alzubi MA, Sohal SS, Olex AL, Dozmorov MG, Harrell JC. Characterizing the efficacy of cancer therapeutics in patient-derived xenograft models of metastatic breast cancer. Breast Cancer Res Treat. 2018 Jul 1;170(2):221–34.

35. Liu Z, Wang Y, Kabraji S, Xie S, Pan P, Liu Z, et al. Improving orthotopic mouse models of patient-derived breast cancer brain metastases by a modified intracarotid injection method. Sci Rep. 2019 Jan 24;9(1):622.

36. Yu EY, Bishop M, Zheng B, Ferguson RM, Khandhar AP, Kemp SJ, et al. Magnetic Particle Imaging: A Novel in Vivo Imaging Platform for Cancer Detection. Nano Lett. 2017 Mar 8;17(3):1648–54.

37. Suzuka H, Mimura A, Inaoka Y, Murase K. Magnetic Nanoparticles in Macrophages and Cancer Cells Exhibit Different Signal Behavior on Magnetic Particle Imaging. J Nanosci Nanotechnol. 2019 Nov 1;19(11):6857–65.

38. Knier NN, Hamilton AM, Foster PJ. Comparing the fate of brain metastatic breast cancer cells in different immune compromised mice with cellular magnetic resonance imaging. Clin Exp Metastasis. 2020 Aug 1;37(4):465–75.

39. Rohani R, de Chickera SN, Willert C, Chen Y, Dekaban GA, Foster PJ. In vivo cellular MRI of dendritic cell migration using micrometer-sized iron oxide (MPIO) particles. Mol Imaging Biol. 2011 Aug;13(4):679–94.

